# Codon usage bias study of the Vitamin D receptor (VDR) gene of Multiple Sclerosis and Diabetes-1 patients

**DOI:** 10.1101/2023.07.06.547960

**Authors:** Sushanta Kumar Barik, Jyotirmayee Turuk, Sidhartha Giri, Sanghamitra Pati

**Author notes:** Address for correspondence: Dr Jyotrimayee Turuk (Scientist-C), ICMR-Regional Medical Research centre-Bhubaneswar, Odisha, India-751023, Email ID.

## Abstract

**Objective:** The codon bias usage study of the 33 Vitamin D Receptor (VDR) genes of the Multiple sclerosis and Diabetes-1 patients were characterized.

**Methods:** Various computational tools such as clustal-W, Codon adaptation index calculation (CAI cal), Effective number of codons (ENC), Relative Synonymous Codon Usage, Codon usage frequency, Nucleotide substation rate calculation, Relative codon deoptimization index, Grand average hydropathicity, Sequence Manipulation Suite software’s were used to find out the codon usage pattern of the VDR genes in both group of patients.

**Results:** The base compositions, nucleotide substation rates, codon adaptation index, hydrophobic nature of the amino acids of VDR gene of both group of patients were analysed. **Conclusion:** The analysis of the synonymous codon usage pattern of the genes would helpful in the heterologous expression of the VDR genes leads to codon optimizations in Multiple Sclerosis and Diabetes-1 patients. The codon usage bias analysis of the VDR gene of the Multiple sclerosis and Diabetes-1 patients through computational approach determined the pattern of VDR gene expression and evolution during the acquiring of the disease in the patients.

## Introduction

Genomic pattern of pleiotropy was observed in organisms. ^[1]^ Many genes in human beings exhibit the phenomenon of pleiotropism. ^[2]^ Human genome genetic variation effects on gene expression that leads to phenotypic variation and disease susceptibility.^[3]^ The vitamin-D receptor gene (VDRG) is around 75kb has pleiotropic effects on biological actions of 1,25-dihydroxy vitamin -D3 to modulate the expression of genes in human beings.^[4]^ 1,25 dihydroxy vitamin D3 (1,25 (OH)2 D3) is a biologically active form of vitamin-D exerts most of its effects on immunosuppressive nature to bind with the specific nuclear receptors. The VDRG polymorphism is associated with the multiple sclerosis in Japanese. ^[5]^ A meta-analysis of the association of the VDR gene polymorphism with the risk of type-1 Diabetes mellitus was reported. ^[6]^

Codon usage bias of a gene has evolved through the genetic drift, mutation, and natural selection in various organisms. Length and expression of the gene, genomic composition, GC content, recombination rates, position, and context of the codon of the genes, gene interaction factors influenced the codon bias activity of the gene in organisms. ^[7]^ Due to the polymorphism in the VDRG gene leads to development of the diseases like Multiple sclerosis and Diabetes-1 in human population, so a codon usage bias analysis was conducted through computational approach.

In the computational study approach, we find out the VDR gene interlinking with the base composition, effective number of codons (ECNS), codon adaptation, relative synonymous codon usage (RSCU), codon frequency measurement, nucleotide substitution rate, relative codon deoptimization index (RSDI), grand average hydropathicity (GRAVY) in patients with Multiple sclerosis and Diabetes-1.

## Methods

The 33VDR gene sequences were extracted from the NCBI website (https://www.ncbi.nlm.nih.gov). The sequences were aligned and taken up for equal length using the clustal W (https://www.ebi.ac.uk/jdispatcher/). The 17 VDR gene sequences lengths ranges from 1-398bp and 16 VDR gene sequences lengths ranges from 1-700bp. The 17 VDR gene sequences ID is available in NCBI: MW331290.1 to MW331307.1 and the 16 VDR gene sequences ID is available in NCBI: KF054040.1 to KF054055.1. The base composition of the nucleotide sequences of the VDR genes were calculated (http://genomes.urv.es/CAIcal). The software was used to calculate the overall nucleotide compositions (A, C, T, and G%), GC content, the nucleotide compositions at the third position (A3, T3, C3, and G3), G + C% at the first (GC1), the second base of codon (GC2), the third base of codon (GC3) of the 360 bp of the 33 VDR gene sequences. The codon usage bias of each gene was also analysed for ‘effective number of codons’ (ENC), which was used by Frank Wright. The ENC ranges from between 20 (an extreme bias as just one codon is used for each amino acid) and 61 (no bias, as all codons are used equally). Effective number of codons (Nc) is a simple measure of the rate of the synonymous codon usage bias in complementary determining sequence of the gene and the number of amino acids independently. Nc can be easily calculated from the codon usage data alone. Nc is a meaningful measure of the extent of the codon preference of a gene. ^[8,9]^ The Nc of the VDR genes could measure by the implementation of the modified formula are available http://agnigarh.tezu.ernet.in/∼ssankar/cub.php. ^[10]^ The codon adaptation index (CAI) of the 33 VDR genes was determined at the E-CAI server (http://genomes.urv.es/CAIcal/E-CAI).^[11]^ The relative synonymous codon usage (RSCU) of the 33 VDR genes was determined at the E-CAI server (http://genomes.urv.es/CAIcal/E-CAI). The codon frequency measurement of the 33 VDR genes will be determined by the help of the sequence manipulation suite (www.bioinformatics.org/sms2/)/. Evolutionary rate of each individual residue of the VDR gene was calculated using MEGA 11. ^[12]^ Relative codon deoptimization index of the 33 VDR genes was calculated (http://genomes.urv.es/CAIcal/E-CAI). Hydropathicity of the amino acids of the 33 VDR genes was calculated (www.bioinformatics.org/sms2/)/. ^[13]^

## Results

### Base composition analysis of the 33 VDR gene

The 17 VDR gene 360 bp nucleotide sequences composition of the multiple sclerosis patients were analysed. The overall nucleotide compositions of 17 VDR gene nucleotide sequences were A (64%), C (116%), T(68%), G(112%). GC content (63.33%, N=14, 63.06%, N= 3), the nucleotide composition at the third position (A3 = 9%, C3=49-50%, T3=14%-15%, G3=47%), G +C% at the first (GC1)=61.67%, GC2=47.50%, GC3= 80.83%. The 16 VDR gene nucleotide composition of type-1 Diabetes patients were calculated and analysed. The overall nucleotide composition of 16 VDR gene of 360bp nucleotide sequences were A(63%), C (113-116), T (68%-70% G (111-114%), the nucleotides of GC content (63.06%N=7, 63.31%, N=2, 63.31%, N=7), the nucleotide composition at the third position (A3 =9%, C3=48-50%, T3=14-15%, G3=47-48%), G+C% at the first (GC1)=61.67%, N=7 ; 62.50%, N=9), GC2=46.67%,N= 1, 47.50%, N=14, 48.33%, N=1, GC3 =79.17%, N=1; 80.00%,N=12; 80.83%, N=3).

### Effective Number of Codon (ENC) calculations of VDR gene patients

ENC can be used to quantify the absolute codon usage bias in coding sequences of 33 VDR gene of Multiple sclerosis and Diabetes patients. The ENC can be implemented through a modified formula are available http://agnigarh.tezu.ernet.in/∼ssankar/cub.php.^[10]^ The relationship between the GC3 value and NC value of each codon of the 33 VDR gene was calculated. ENC of 17 VDR gene sequences of Multiple Sclerosis Patients are given in table-1 and ENC of 16 VDR gene sequences of Diabetes-1 patients are given in table-2.

### Codon adaptation index (CAI) analysis of VDR gene

Codon adaptation index (CAI) is a simple effective measure in degree of codon usage bias towards the major codons. It can be calculated by comparing its codon usage frequency with a reference set of highly expressed genes from a species. CAI score for a gene is calculated from the frequency of all codons in a gene where the value ranges between 0.0 and 1.0. Higher CAI value means a stronger codon usage bias and a higher expression level (Salim et al, 2008). The average eCAI value of the 17 VDR gene of Multiple sclerosis patients was 0.81. The VDR genes of multiple sclerosis patients shown the moderate level of expression as per the above eCAI value. The average eCAI value of the 16 VDR gene of Diabetes patients was 0.81 and this value shown the moderate level of expression of VDR gene in Diabetes patients.

### Relative synonymous codon usage analysis

The frequent distribution codon of each isolate information is solved by RSCU. The value of RSCU 1 indicates that the codon is used as expected by random usage, RSCU of >1 indicates a codon used more frequently than expected randomly but <1 indicates a codon is used less frequently than random. The pattern of relative synonymous codon usage (RSCU) of 17 VDR gene of Multiple sclerosis patient was calculated and the value of each codon was presented. The pattern of RSCU of 17 VDR gene of Multiple sclerosis patient is given below:

1. RCSU of the VDR gene of Multiple sclerosis

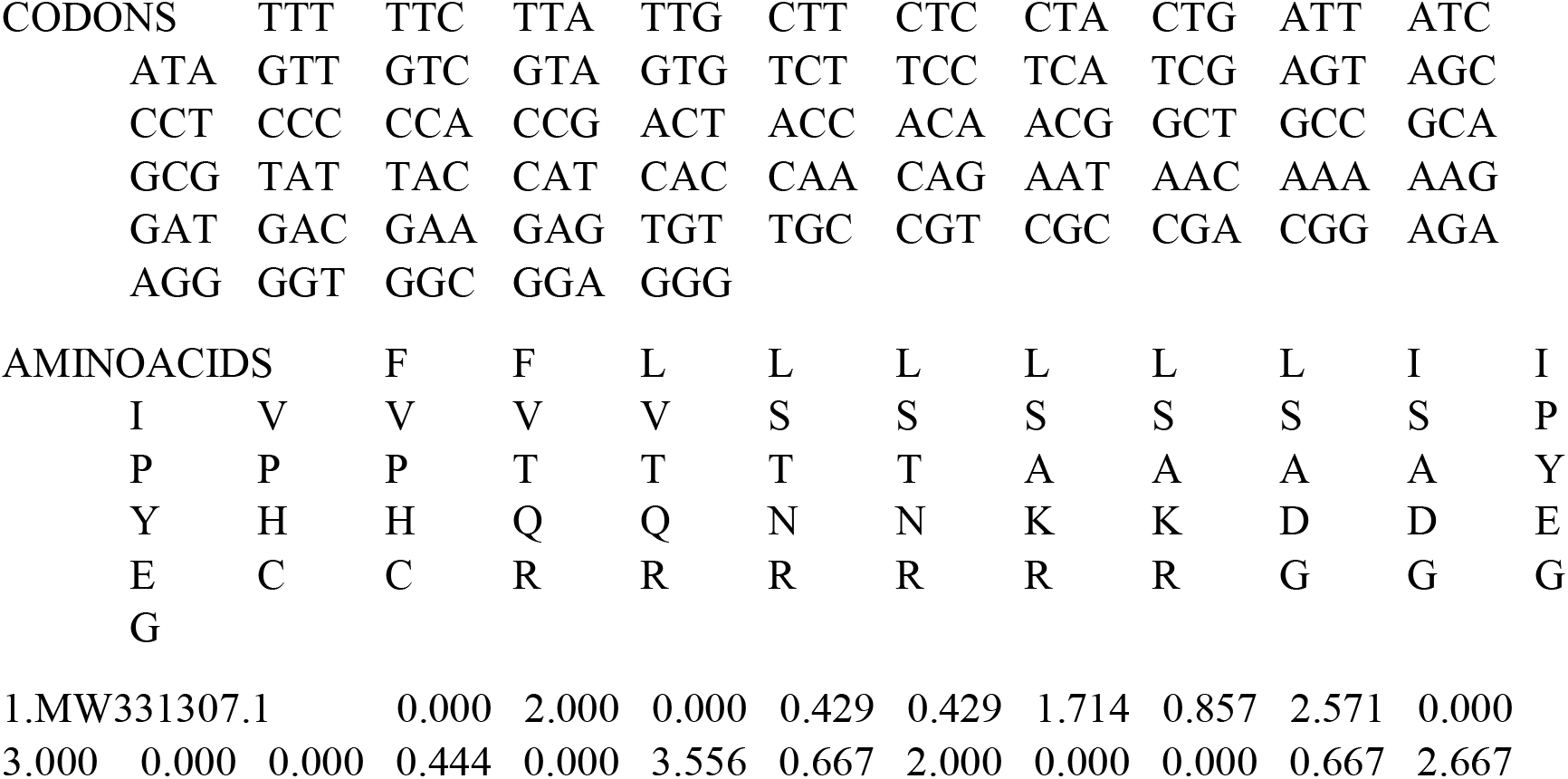
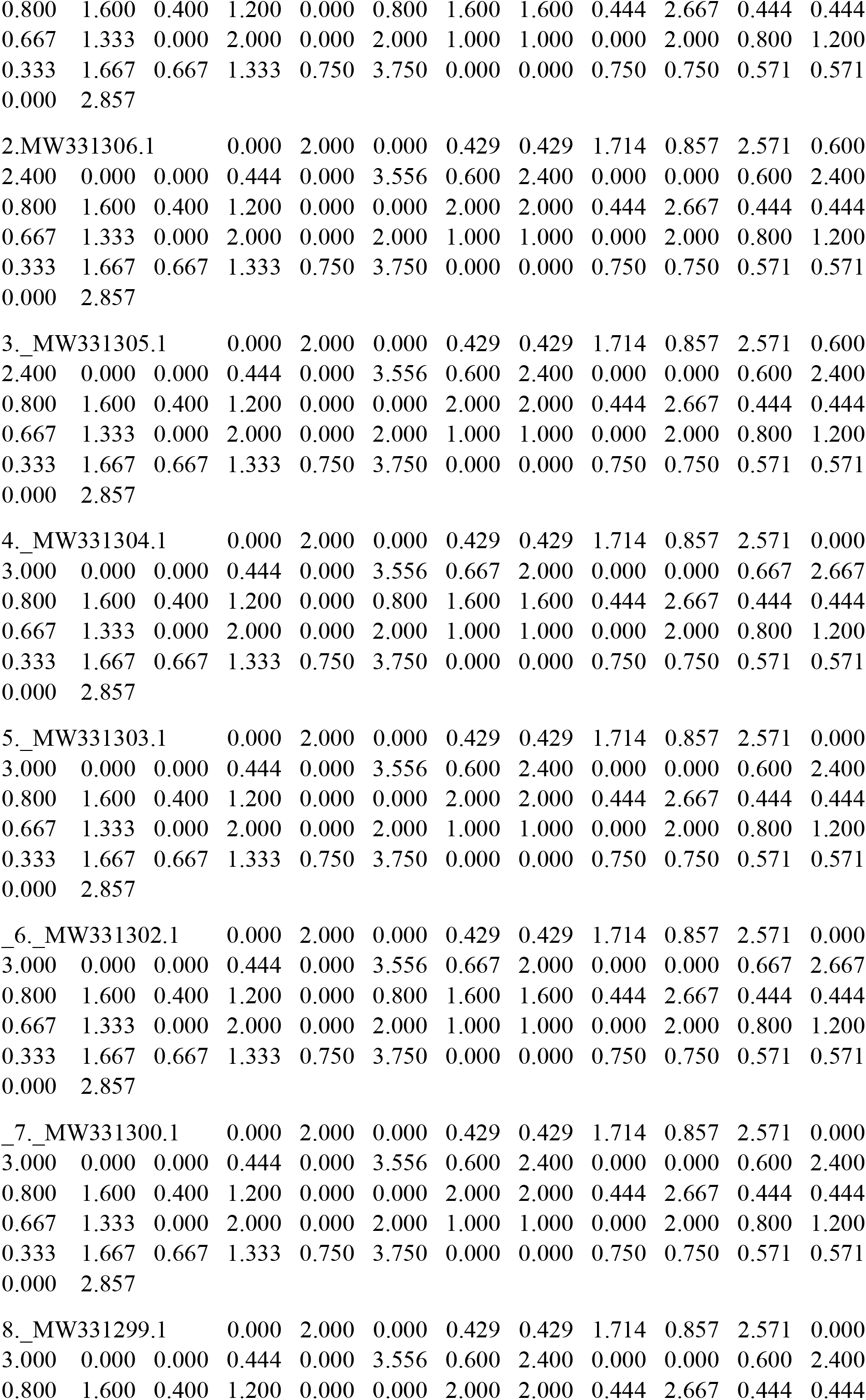
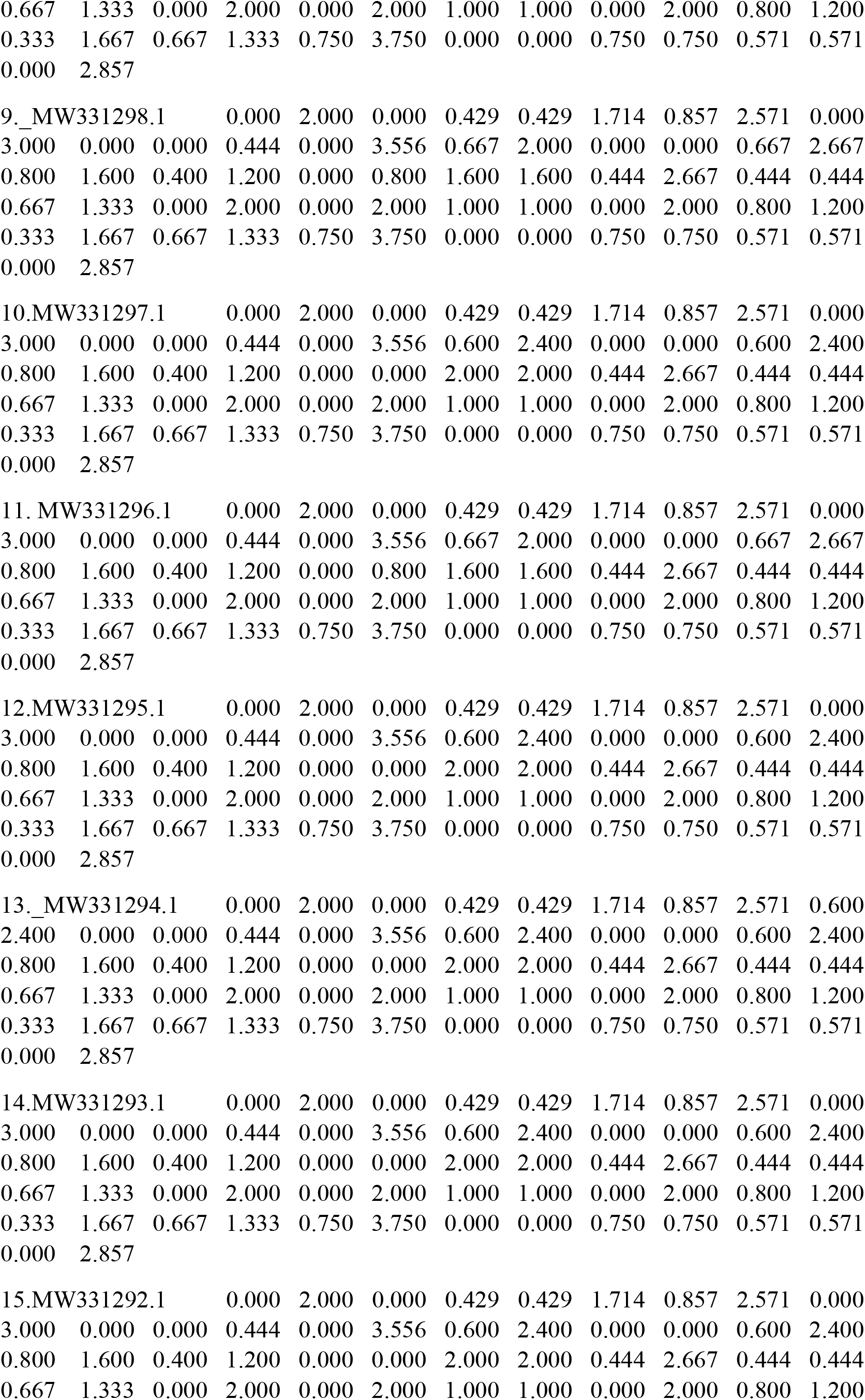
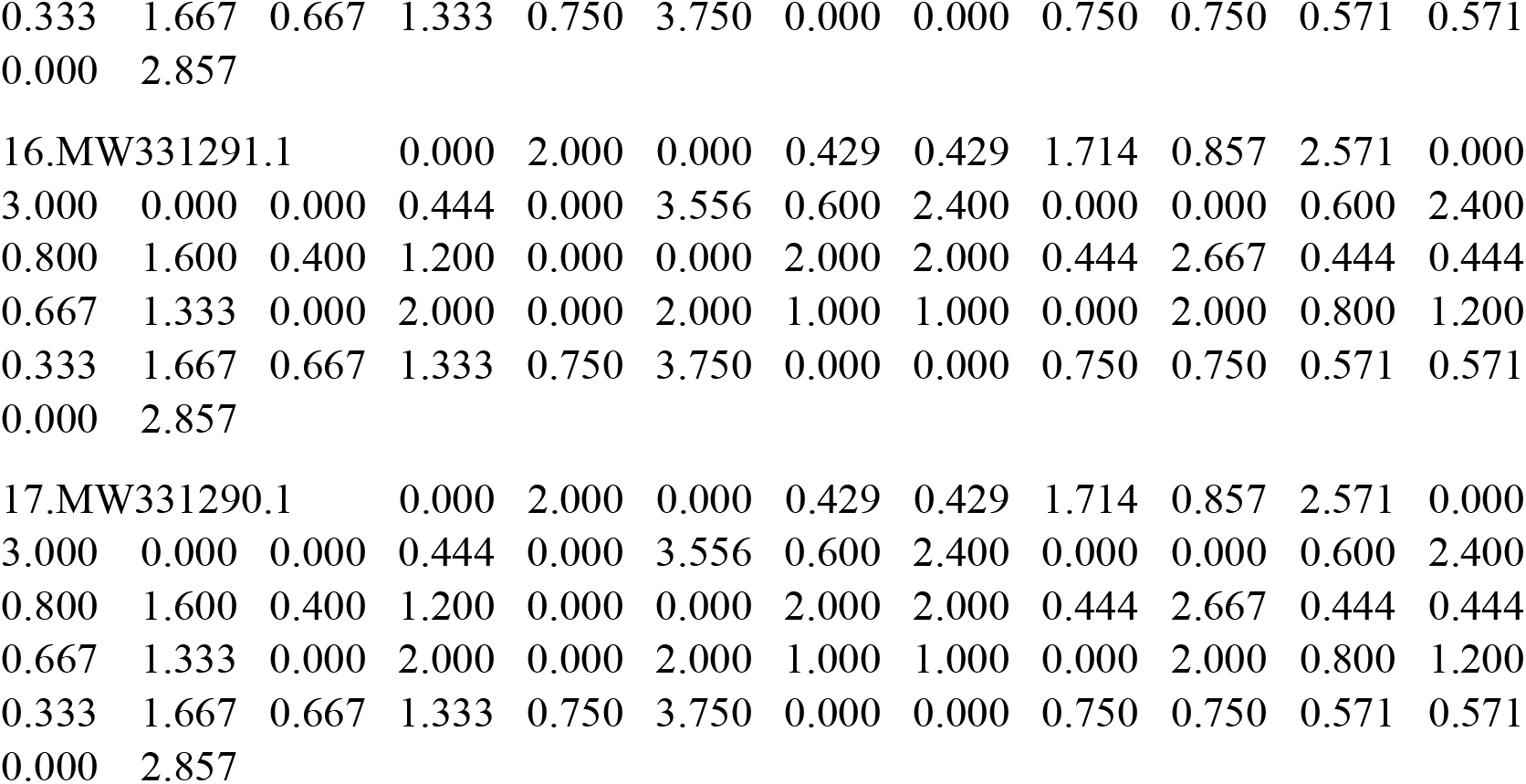

The pattern of RSCU of 16 VDR gene of Diabetes patient is given below:

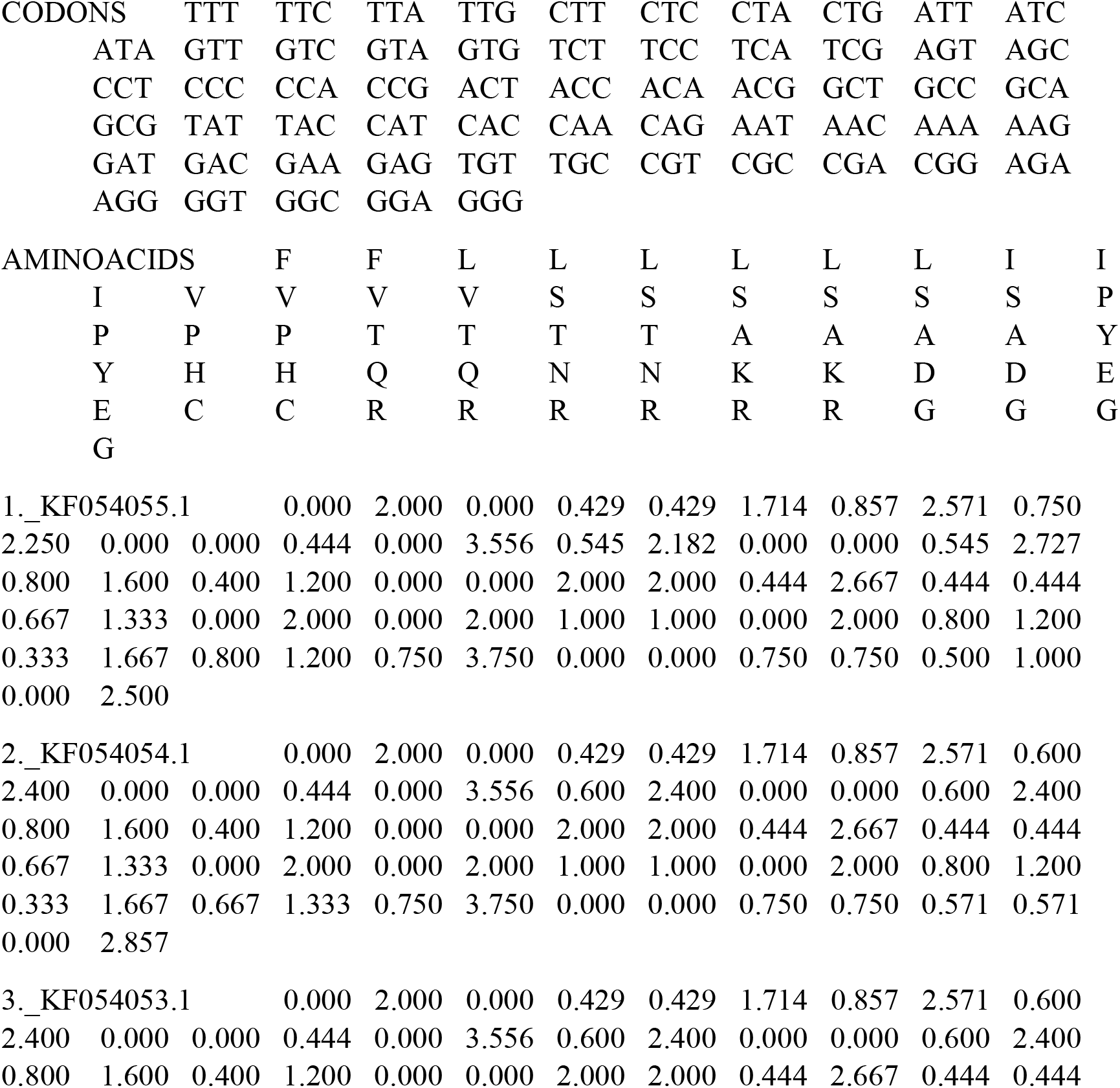
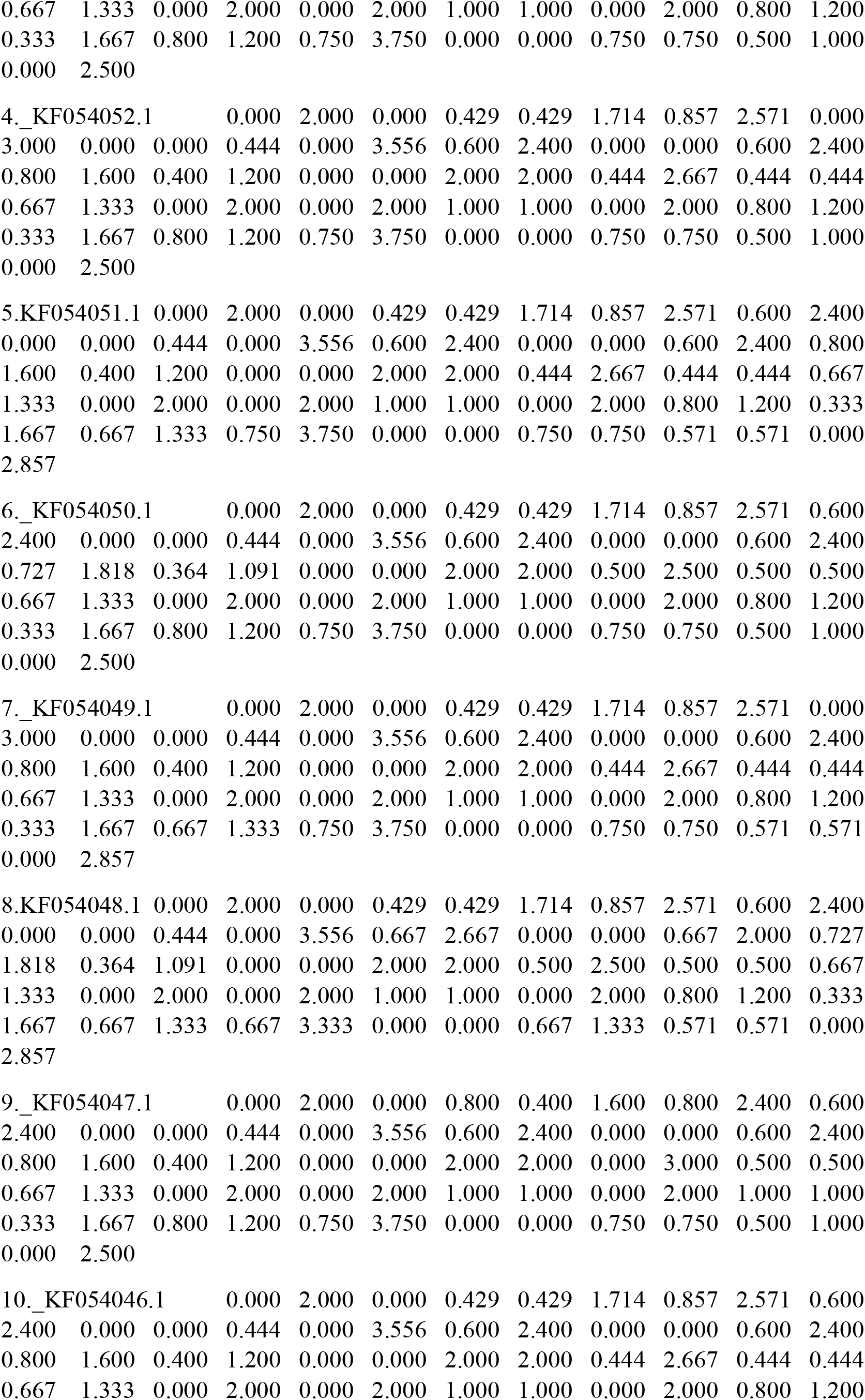
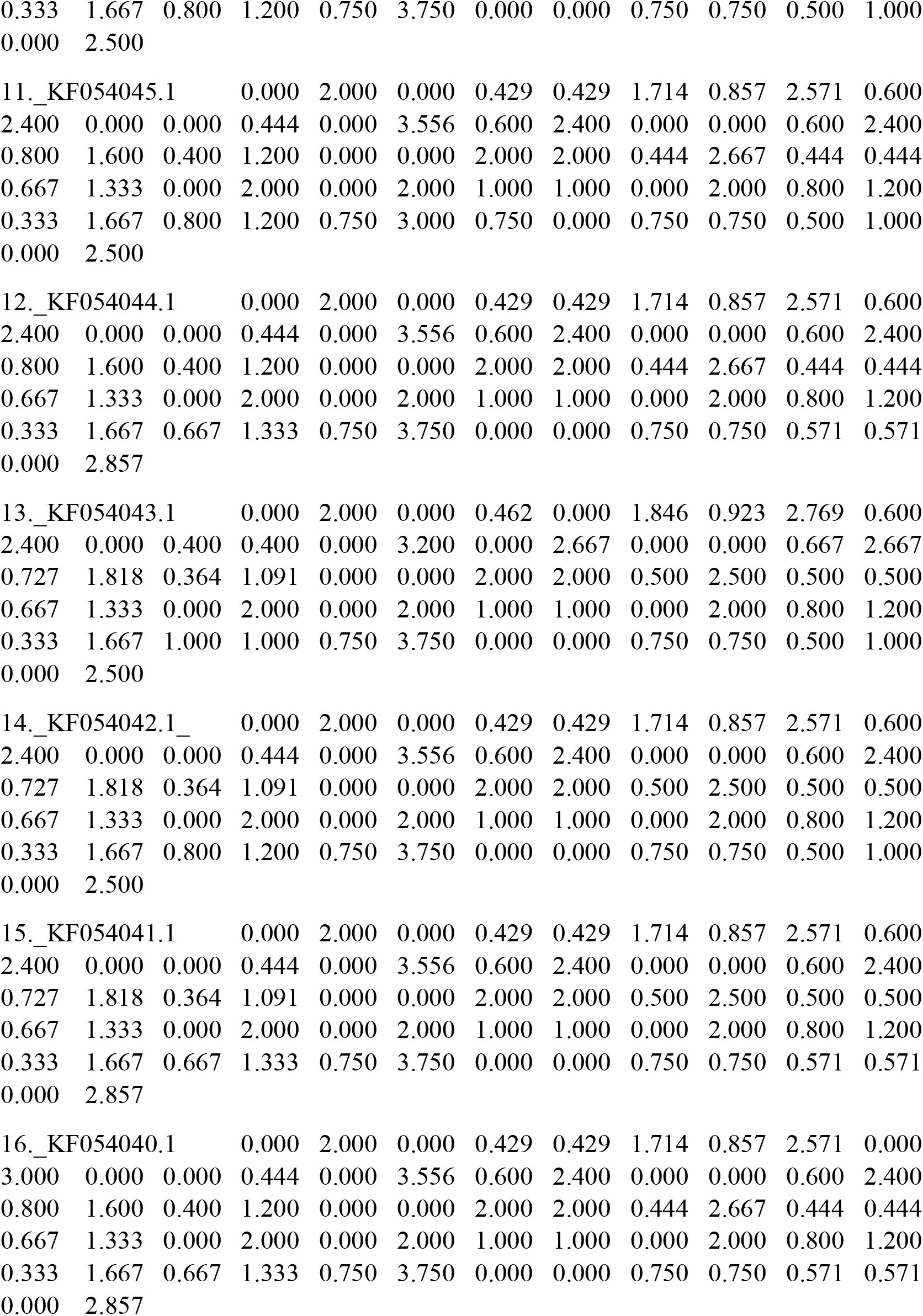

### Codon usage frequency analysis

Codon usage frequency analysis accepts one or more DNA sequences to analyse the VDR gene sequences to returns the number and frequency of each codon and find out the frequency of each codon that code for the same amino acids (Synonymous codons). The synonymous codons of 17 VDR gene of Multiple sclerosis patients were analysed and 16 VDR gene of Diabetes patients were analysed. The synonymous codons analysis of the 17 VDR gene of Multiple sclerosis patients are given in the supplementary file-1 and 16 VDR gene of Diabetes patients are given in the supplementary file-2.

### Nucleotide substitution rate calculation

The nucleotide substation rate was calculated based on Maximum Likelihood Estimate of Substitution matrix. Substitution pattern and rates were estimated under Tamura-Nei (1993) model. 17 VDR gene nucleotide substitutions rates of Multiple sclerosis patients were calculated and given in the table-3. The 16 VDR gene nucleotide substation rates were calculated and given in the table-4.

### Relative codon deoptimization index (RCDI)

The relative codon deoptimization index (RCDI) values for the 17 VDR gene of multiple sclerosis patients and 16 VDR gene of Diabetes patients were calculated using the RCDI/eRCDI server. The RCDI value 1 indicates that virus adopts to the host codon usage pattern and displays a host codon adapted usage pattern. The RCDI values higher than 1 indicate the deoptimization of the codon usage pattern of the virus within the host. ^[14]^ The RCDI/eRCDI value of 17 VDR gene of Multiple sclerosis patients was 1.83/2.21. The RCDI/eRCDI value of the 16 VDR gene of Diabetes patients was 1.77/2.11.

### Grand average hydropathicity (GRAVY)

Grand average hydropathicity of the protein sequence was calculated by adding the hydropathy value of each residue of a protein and its length. ^[15]^ Grand average hydropathy (GRAVY) of each protein sequences of 17 VDR gene of Multiple sclerosis patients and 16 VDR gene of Diabetes patients were calculated. The value of GRAVY of 17 VDR gene of Multiple sclerosis patients were 0853 (N=3), 0.862 (N=9),0.869 (N=5). The GRAVY of 16 VDR gene of 16 Diabetes patients were 0.844 (1), 0.846 (1), 0.852 (1), 0.853(4), 0.854 (2), 0.855(1), 0.861(1), 0.862 (4), 0.863(1).

## Discussion

The codon and amino acid usage shaped the genomic structure and evolutionary pattern of organisms. The GCsyn content of amino acids were corelates with the codon/amino acid usage. The GC content determined the usage variation of amino acids in a gene. ^[16]^ The GC content of the VDR gene of both type-1 Diabetes and Multiple sclerosis patients were observed to be high. Effective number of codon calculation investigated the codon bias shaped of the gene of organisms and participate the major role of evolutionary trend of the gene of the organisms. The variation within the genome of the codon usage bias is driven by the codon selection forces.^[17]^ The GC content of VDR gene of the Multiple sclerosis and Diabetes -1 patients was more than 63% even the improved effective number of codons were more biased due to the codon selection forces of the genes of the patients. Codon bias usage (CBU) of the gene is the cause of the degeneracy of the genetic code that uses multiple synonymous codons code for the same amino acid. The CAI value is associated with the virus-host interaction in the molecular evolution. CAI value of the gene revealed the less adaptation in the host-virus interaction.^[18]^ The relative synonymous codon adaptation index (RCAI) is a sensitive measure in codon usage bias study. The RCAI discriminate between the highly biased and unbiased regions of the gene of the organisms.^[19]^ Codon usage differs in all genes of the genome of an organism. CAI value of the certain organism is based on the codon usage frequency of the highly expressed genes and find out the expression level of the gene of an organism. ^[20]^ From the analysis of the VDR genes of the Multiple sclerosis and Diabtetes-1 patients indicates the moderate level of expression of the gene of an organism. The RSCU analysis of the pancreatitis associated genes confirmed the synonymous and non-synonymous codon variants of the gene is associated with the disease initiation and progression. ^[21]^ 59 codons are exhibits as synonymous codon in 33 VDR gene of Multiple sclerosis patients and Diabetes-1 patients. The synonymous codons were characterized in the HPRT1 gene of the Mammalian species. ^[22]^ In our analysis, a greater number of synonymous codons are observed in the VDR gene of Multiple sclerosis and Diabetes-1 patients. It RSCU results indicates, the existence of synonymous codons in the VDR gene of Multiple Sclerosis and Diabetes-1 patients. Codon usage bias reflects the genes of genome between the mutational biases and natural selection of the organisms. ^[23]^ Trend between the usage of amino acids and its occurrence of the gene is observed by the analysis of the Codon usage frequency of amino acids in the genome of organisms. The analysis of the standard genetic code realized in the development of modern life and the evolvement of genetic code in the development of normal life.^[24]^ The codon frequency analysis of the VDR gene of the Multiple sclerosis and Diabtetes-1 patients indicates the communication of life of human beings during the progression of the disease. The surprising changes in the protein and DNA sequences of the genome of the organisms brought a theoretical understanding of the neutral mutations and genetic drifts as well as the substitution rate of mutations towards genomic evolution of the organisms.^[25]^ The evolutionary pressure influenced both the codon bias and translation initiation efficiency of highly expressed genes of an Organism. ^[26]^ The nucleotide substation rate of the VDR gene of Multiple sclerosis and Diabetes-1 patients was different due to variation in the number of synonymous codons. The optimal codons of the genes of Novel porcine parvoviruses-2 were frequently reported through RCDI. ^[27]^ The RCDI value of VDR genes of the Multiple sclerosis and Diabetes -1 patients were slightly different and suggesting the evolution of the optimal codons in the genome of the organisms.

Thus, the computational analysis of the codon bias usage genomic signatures may help to find out suitable drugs to treat the Multiple sclerosis and Diabetes-1 patients in vitamin -D deficiency in the field of genomic medicine.

## Conclusion

Natural selection and genomic drift in the genes of the genome of the organisms is a selection pressure phenomenon leads to molecular evolution. The codon usage bias analysis of the VDR gene of the Multiple sclerosis and Diabetes-1 patients using the computational approach would suggest the VDR gene expression and evolution during the acquiring of the disease of the patients.

Retrieved VDR gene sequences of Multiple Sclerosis and Diabetes-1 patients’ number from NCBI, USA: Gene ID of Multiple Sclerosis: MW331307.1, MW331306.1, MW331305.1, MW331304.1, MW331303.1, MW331302.1, MW331300.1, MW331299.1, MW331298.1, MW331297.1, MW331296.1, MW331295.1, MW331294.1, MW331293.1, MW331292.1, MW331291.1, MW331290.1. Gene ID of Diabetes-1 patients: KF054055.1, KF054054.1, KF054053.1, KF054052.1, KF054051.1, KF054050.1, KF054049.1, KF054048.1, KF054047.1, KF054046.1, KF054045.1, KF054044.1, KF054043.1, KF054042.1, KF054041.1, KF054040.1.

## Declaration statements

### Competing interest

All authors are declared no conflicting financial interest.

### Funding Statement

Indian Council of Medical Research, Govt. of India for providing ICMR-RA fellowship (file no:3/1/2/299/2021-Nut) to Dr Sushanta Kumar Barik.

### Authorship contribution statement

SKB: Analyse the VDR gene sequences through various software’s, Write and Review the manuscript, JT: Write and Review the Manuscript, SG: Review the Manuscript, SP: Review the manuscript.

## Acknowledgement

ICMR, Govt. of India is acknowledged for the ICMR-RA fellowship.

## Preprint

Codon usage bias study of the Vitamin D receptor (VDR) gene of Multiple Sclerosis and Diabetes-1 patients. doi: https://doi.org/10.1101/2023.07.06.547960.

## Notes

### Competing Interest Statement

The authors have declared no competing interest.

### Summary of Updates

There is no significant difference between this version and previous version of the manuscript.

